# Chromatin modifiers alter recombination between divergent DNA sequences

**DOI:** 10.1101/519397

**Authors:** Ujani Chakraborty, Beata Mackenroth, David Shalloway, Eric Alani

**Author notes:** Corresponding Author: Eric Alani, Ph.D., Cornell University Department of Molecular Biology and Genetics, 459 Biotechnology Building, Ithaca, NY 14853-2703, United States of America.

## Abstract

Recombination between divergent DNA sequences is actively prevented by heteroduplex rejection mechanisms. In baker’s yeast such anti-recombination mechanisms can be initiated by the recognition of DNA mismatches in heteroduplex DNA by MSH proteins, followed by recruitment of the Sgs1-Top3-Rmi1 helicase-topoisomerase complex to unwind the recombination intermediate. We previously showed that the repair/rejection decision during single-strand annealing recombination is temporally regulated by MSH protein levels and by factors that excise non-homologous single-stranded tails. These observations, coupled with recent studies indicating that mismatch repair factors interact with components of the histone chaperone machinery, encouraged us to explore roles for epigenetic factors and chromatin conformation in regulating the decision to reject vs. repair recombination between divergent DNA substrates. This work involved the use of an inverted repeat recombination assay thought to measure sister chromatid repair during DNA replication. Our observations are consistent with the histone chaperones CAF-1 and Rtt106 and the histone deacetylase Sir2 acting to suppress heteroduplex rejection and the Rpd3, Hst3 and Hst4 deacetylases acting to promote heteroduplex rejection. These observations and double mutant analysis have led to a model in which nucleosomes located at DNA lesions stabilize recombination intermediates and compete with mismatch repair factors that mediate heteroduplex rejection.

**Summary:** Recombination between divergent DNA sequences is actively prevented by heteroduplex rejection mechanisms. In this study we explored roles for epigenetic factors and chromatin conformation in regulating the decision to reject vs. repair recombination between divergent DNA substrates. Our observations are consistent with the histone chaperones CAF-1 and Rtt106 and the histone deacetylase Sir2 acting to suppress heteroduplex rejection and the Rpd3, Hst3 and Hst4 deacetylases acting to promote heteroduplex rejection. These observations have led to a model in which nucleosomes located at DNA lesions stabilize recombination intermediates and compete with mismatch repair factors that mediate heteroduplex rejection.

## Introduction

Homologous recombination (HR) is a conservative DNA repair pathway that is critical for repairing DNA double strand breaks (DSBs). This process is regulated to prevent recombination between divergent DNA sequences (reviewed in George and Alani 2012). Such regulation, which can prevent deleterious chromosomal rearrangements, is initiated through the recognition of mismatches in heteroduplex DNA that form during strand invasion steps involving single-stranded DNA from a broken chromosome and a divergent duplex donor. In the yeast *Saccharomyces cerevisiae*, the Msh2-Msh6 and Msh2-Msh3 mismatch repair (MMR) complexes recognize mismatches in heteroduplex DNA and recruit the RecQ family helicase-topoisomerase complex Sgs1-Top3-Rmi1 to unwind the recombination intermediate in a process known as heteroduplex rejection (Datta *et al*. 1996; Chen and Jinks-Robertson 1999; Nicholson *et al*. 2000; Myung *et al*. 2001; Spell and Jinks-Robertson 2004; Sugawara *et al*. 2004; Goldfarb and Alani 2005; Chakraborty *et al*. 2016; Hum and Jinks-Robertson 2019). If rejection does not occur, MSH complexes initiate repair of the mismatches in heteroduplex DNA after/while the break is repaired. We refer to this regulation as the repair/rejection decision.

An important aspect of the repair/rejection decision is that tolerating multiple mismatches in heteroduplex DNA formed from divergent substrates can lead to chromosomal rearrangements, but a highly efficient rejection system can result in DSBs that are not repaired. Various factors are likely to influence this decision (reviewed in Chakraborty and Alani 2016). For example, we found that MSH protein levels influence the repair/rejection decision during single strand annealing (SSA; Chakraborty *et al*. 2016). During SSA, HR is initiated by a DSB located between two closely spaced repeat sequences. Resection of the DSB promotes annealing of homologous sequences, followed by the clipping of 3’ non-homologous tails that must be excised before repair steps is completed. During SSA involving divergent repeat sequences, modest overexpression of Msh6 resulted in a significant increase in heteroduplex rejection due to a decrease in the availability of Msh2-Msh3 to clip the 3’ tails. Thus 3’ tail clipping during SSA serves as a regulatory step, with rejection favored prior to 3’ tail removal. Consistent with these findings, Anand *et al.* (2017) showed in a break-induced repair recombination system that 3’ non-homologous tails promote heteroduplex rejection, and the absence of such tails prevents it. These observations indicate a crosstalk between the rejection machinery and the local environment that likely occurs prior to steps in HR that involve repair DNA synthesis.

Recent studies have indicated that chromatin structure can influence HR outcomes. Several nucleosome remodeling complexes have been shown to be recruited to DSBs in steps thought to increase chromatin accessibility and allow strand resection and presynaptic filament formation (reviewed in Hauer and Gasser 2017). Such chromatin remodelers were shown to promote chromatin mobility during DSB formation, and the increased mobility correlated to more efficient repair by HR (Dion *et al*. 2012; Mine-Hattab and Rothstein 2012; Neumann *et al*. 2012; Hauer *et al.* 2017). Also, histone chaperones, which act in DNA replication-dependent nucleosome assembly, have been implicated in DNA repair. These chaperones include CAF-1, Asf1 and Rtt106, which are all involved in DNA replication-dependent nucleosome assembly (Tyler *et al.* 1999; Tagami *et al*. 2004; Huang *et al*. 2005). For example, nucleosome assembly mediated by CAF-1 during DNA repair is coupled with DNA synthesis and requires an interaction with PCNA (Gaillard *et al*. 1996; Tyler *et al*. 1999; Linger and Tyler 2005; Polo, Roche and Almouzni 2006; Pietrobon *et al*. 2014), and CAF-1 and Asf1 play roles in restoring chromatin after DNA repair in budding yeast at repair sites by turning off the DNA damage checkpoint (Chen *et al*. 2008; Kim and Haber 2009; Diao *et al.* 2017).

Histone modifications have also been shown to affect genome stability. For example, deacetylation of an acetylated lysine residue at amino acid 56 in Histone 3 (H3K56) by Hst3 and Hst4 is required for the suppression of mutations and gross chromosomal rearrangements, and acetylation of H3K56 by Rtt109 is also required for suppression of mutations (Kadyrova *et al*. 2013). In contrast, histone deacetylases such as Rpd3L and Hda1 promote trinucleotide repeat expansions associated with various neurodegenerative diseases (Debacker *et al*. 2012). Thus, histone acetylation and deacetylation are likely to play important roles in various stages of HR. In support of roles for acetylases and deacetylases in HR, Tamburini and Tyler (2005) showed that histone acetyltransferases such as Gcn5 and Esa1 are recruited to an HO endonuclease induced DSB in *S. cerevisiae* followed at a later stage by recruitment of histone deacetylases such as Sir2, Hst1 and Rpd3. Furthermore, they showed that mutating acetylable lysine residues in Histone subunit 4, or deleting *GCN5* or *RPD3,* caused inviability in response to HO endonuclease induced lesions repaired primarily by HR. These observations suggest that after DSB formation, histone acetylases participate in nucleosome removal in the vicinity of the break and “open up” the chromatin structure increasing the accessibility of the underlying DNA to repair factors. Once DNA repair is underway, histone deacetylases likely modify the chromatin to a “closed” conformation that serves as a commitment step to complete the repair process and restore chromatin to its original state.

The NAD-dependent histone deacetylase Sir2, known for its role in transcriptional silencing and heterochromatin formation, also plays direct roles in forming a repressive local chromatin environment around most euchromatic replication origins (Gartenberg and Smith 2016; Hoggard *et al*. 2018). Sir2 is also involved in DNA repair pathways such as non-homologous end joining and nucleotide excision repair (Tsukamoto, Kato, and Ikeda 1997; Boulton and Jackson 1998; Guintini *et al*. 2017). *sir2* mutants are hypersensitive to DNA damaging agents, and a number of studies have reported that in response to DNA DSBs, a significant fraction of the histone bound SIR complex was displaced from subtelomeric regions and relocated to sites of DSBs in a DNA checkpoint dependent manner, suggesting that the recruitment of SIR complexes reflects the assembly of a repressed chromatin state following DNA repair (Martin *et al*. 1999; McAinsh *et al*. 1999; Mills, Sinclair, and Guarente1999).

A number of studies have shown that nucleosome assembly on newly synthesized DNA during replication and MMR are mutually inhibitory processes. MMR during DNA replication is thought to be restricted to the short time window between the formation of the mismatch and the chaperone-assisted assembly of nucleosomes on the newly replicated DNA (Li *et al*. 2009; Kadyrova, Blanko and Kadyrov 2011; Schopf *et al*. 2012; Blanko, Kadyrova and Kadyrov 2016). The human MSH2-MSH6 complex has been shown to interact with CAF-1 *in vitro* (Schopf *et al.* 2012). Additionally, human MSH2-MSH6 inhibits CAF-1 and ASF1A-dependent packaging of a DNA mismatch into a nucleosome, and deposition of the (H3-H4)_2_ tetramers on DNA protects the discontinuous daughter strand from unnecessary degradation during MMR (Blanko, Kadyrova and Kadyrov 2016). Moreover, Pietrobon *et al*. (2014) showed in fission yeast that CAF-1 stabilizes D-loops in an HR pathway, by counteracting their disassembly mediated by the RecQ family helicase Rqh1 when cells replicate a damaged template.

How is the balance between heteroduplex rejection and homologous recombination repair maintained? We tested in baker’s yeast roles for histone acetylases, deacetylases, and chaperones in regulating the repair/rejection decision. To our knowledge, this is the first study to examine roles for epigenetic factors and chromatin modifiers in modulating heteroduplex rejection. We show that histone chaperones CAF-1 and Rtt106 suppress heteroduplex rejection in steps dependent on mismatch recognition. Additionally, the histone deacetylase Sir2 appears to act in a common pathway with CAF-1 and/or Rtt106 to suppress rejection. However, other factors involved in nucleosome assembly during DNA replication such as Asf1 and Rtt109 do not affect rejection efficiency. Similarly, histone acetylases such as Gcn5 that assemble early at recombination sites, and other histone deacetylases such as Hst1 do not affect the efficiency of rejection. However, mutants lacking the Hst3, Hst4, or Rpd3 histone deacetylases show defects in rejection. Taken together, these results are consistent with the idea that nucleosomes at DNA lesions, which are likely to be localized independently of DNA synthesis, stabilize recombination intermediates and thus prevent access/unwinding by anti-recombination factors.

## Materials and Methods

### Yeast strains and plasmids

Yeast strains used in this study are listed in Table 1 and were constructed and grown using standard techniques (Rose, Winston, and Hieter 1990). Gene disruptions used to make the strains are described in Tables 1 and 2 (*geneXΔ∷KANMX*) and were obtained by PCR amplification (details provided upon request) of chromosomal DNA derived from the yeast knockout collection (Brachmann *et al.* 1998). PCR products, linear DNA fragments obtained from pEAI98 (*msh2Δ∷hisG-URA3-hisG)* and pEAA633 (*pol30-8∷KANMX*, see below), and *2μ* vectors (see below) were introduced into yeast strains using standard transformation procedures (Gietz and Schiestl 1991). The presence of mutant alleles was confirmed by PCR analysis of chromosomal DNA, and in some cases DNA sequencing of the amplified DNA fragments.

**Table 1.** Strains used in this study.

|  |  |
| --- | --- |
| SJR328 | *MATα*, *ade2-101*, *his3Δ200*, *ura3-Nhe*, *lys2ΔRV::hisG*, *leu2-R* |
| SJR769 | SJR328, *cβ2/cβ2::LEU2*, homologous 350 bp substrates |
| GCY615 | SJR328, *cβ2/cβ2-ns::LEU2*, predicted to form four, base-base mismatches in the 350 bp *cβ2* substrates |
| GCY559 | SJR328, *cβ2/cβ2-4L::LEU2*, predicted to form four 4-nt loops in the 350 bp *cβ2* substrates |
| EAY1605,1606 | SJR769, *msh2Δ::hisG-URA3-hisG* |
| EAY1609,1610 | GCY615, *msh2Δ::hisG-URA3-hisG* |
| EAY3853-3856 | SJR769, *hst3Δ::KANMX* |
| EAY3857-3860 | GCY615, *hst3Δ::KANMX* |
| EAY3865-3868 | SJR769, *hst4Δ::KANMX* |
| EAY3869-3872 | GCY615, *hst4Δ::KANMX* |
| EAY3877-3880 | SJR769, *hst3Δ::KANMX hst4Δ::NATMX* |
| EAY3881-3884 | GCY615, *hst3Δ::KANMX hst4Δ::NATMX* |
| EAY3964-3967 | SJR769, *hst1Δ::KANMX* |
| EAY3968-3971 | GCY615, *hst1Δ::KANMX* |
| EAY3841-3844 | SJR769, *sir2Δ::KANMX* |
| EAY3845-3848 | GCY615, *sir2Δ::KANMX* |
| EAY3849-3852 | GCY559, *sir2Δ::KANMX* |
| EAY4002-4005 | SJR769, *rpd3Δ::KANMX* |
| EAY4006-4009 | GCY615, *rpd3Δ::KANMX* |
| EAY4047-4050 | SJR769, *gcn5Δ::KANMX* |
| EAY4051-4054 | GCY615, *gcn5Δ::KANMX* |
| EAY3889-3892 | SJR769, *cac1Δ::KANMX* |
| EAY3893-3896 | GCY615, *cac1Δ::KANMX* |
| EAY3897-3900 | GCY559, *cac1Δ::KANMX* |
| EAY3972,3974,3975 | SJR769, *rtt106Δ::KANMX* |
| EAY3976-3978 | GCY615, *rtt106Δ::KANMX* |
| EAY3901-3904 | SJR769, *asf1Δ::KANMX* |
| EAY3905-3908 | GCY615, *asf1Δ::KANMX* |
| EAY3909-3912 | GCY559, *asf1Δ::KANMX* |
| EAY4127-4129 | SJR769, *pol30-8::KANMX* |
| EAY4130-4132 | GCY615, *pol30-8::KANMX* |
| EAY3925, 3926 | SJR769, *cac1Δ::KANMX asf1Δ::NATMX* |
| EAY3929, 3930 | GCY615, *cac1Δ::KANMX asf1Δ::NATMX* |
| EAY3933, 3934 | GCY559, *cac1Δ::KANMX asf1Δ::NATMX* |
| EAY4018-4021 | SJR769, *rtt106Δ::KANMX asf1Δ::NATMX* |
| EAY4022-4025 | GCY615, *rtt106Δ::KANMX asf1Δ::NATMX* |
| EAY3913-3916 | SJR769, *rtt109Δ::KANMX* |
| EAY3917-3920 | GCY615, *rtt109Δ::KANMX* |
| EAY4010-4013 | SJR769, *rtt106Δ::KANMX cac1Δ::NATMX* |
| EAY4014-4017 | GCY615, *rtt106Δ::KANMX cac1Δ::NATMX* |
| EAY4027-4029 | SJR769, *sir2Δ::KANMX cac1Δ::NATMX* |
| EAY4030-4033 | GCY615, *sir2Δ::KANMX cac1Δ::NATMX* |
| EAY4034-4037 | SJR769, *rtt106Δ::KANMX sir2Δ::NATMX* |
| EAY4038-4041 | GCY615, *rtt106Δ::KANMX sir2Δ::NATMX* |
| EAY4042-4044 | SJR769, *rtt106Δ::KANMX cac1Δ::NATMX sir2Δ::HYGMX* |
| EAY4045,4046 | GCY615, *rtt106Δ::KANMX cac1Δ::NATMX sir2Δ::HYGMX* |
| EAY4069-4072 | SJR769, *cac1Δ::KANMX msh2Δ::hisG-URA3-hisG* |
| EAY4073-4076 | GCY615, *cac1Δ::KANMX msh2Δ::hisG-URA3-hisG* |
| EAY4077-4080 | SJR769, *rtt106Δ::KANMX msh2Δ::hisG-URA3-hisG* |
| EAY4081-4084 | GCY615, *rtt106Δ::KANMX msh2Δ::hisG-URA3-hisG* |
| EAY4216-4218 | SJR769, *sir2Δ::KANMX msh2Δ::hisG-URA3-hisG* |
| EAY4220,4222,4223 | GCY615, *sir2Δ::KANMX msh2Δ::hisG-URA3-hisG* |

SJR328, SJR769, GCY615, and GCY559 were obtained from Sue Jinks-Robertson and are described in Nicholson *et al.* (2000).

**Table 2.** Recombination rates as measured in the inverted repeat reporter assay.

| Genotype | Cβ2/Cβ2 His<sup>+</sup> homologous recombination rate (x10<sup>-6</sup>) | Cβ2/Cβ2-ns His<sup>+</sup> divergent recombination rate (x10<sup>-6</sup>) | Cβ2/Cβ2 rate/<br>Cβ2/Cβ2-ns rate<sup>a</sup> |
| --- | --- | --- | --- |
| wild-type | 0.52 (0.48-0.56)<sup>b</sup> | 0.027 (0.024-0.029)<sup>b</sup> | 20 (17-22)<sup>b</sup> |
| *msh2*Δ | 0.92 (0.70-1.15) | 2.55 (2.31-2.80) | 0.36 (0.28-0.47) |
| *cac1*Δ | 1.19 (1.00-1.38) | 0.030 (0.021-0.040) | 40 (28-57) |
| *rtt106*Δ | 1.87 (1.55-2.20) | 0.028 (0.018-0.038) | 68 (46-101) |
| *asf1*Δ | 3.96 (3.30-4.61) | 0.27 (0.22-0.32) | 15 (11-19) |
| *rtt109*Δ | 3.59 (2.93-4.24) | 0.22 (0.16-0.27) | 16 (12-23) |
| *cac1*Δ *asf1*Δ | 0.60 (0.32-0.89) | 0.093 (0.059-0.13) | 6.4 (3.5-12) |
| *rtt106*Δ *asf1*Δ | 1.98 (1.65-2.31) | 0.14 (0.097-0.18) | 14 (10-20) |
| *cac1*Δ *rtt106*Δ | 2.33 (1.85-2.80) | 0.018 (0.0095-0.027) | 134 (78-229) |
| *pol30-8* | 0.74 (0.61-0.87) | 0.026 (0.020-0.033) | 28 (21-39) |
| *hst3*Δ | 1.67 (1.44-1.89) | 0.090 (0.066-0.11) | 19 (14-25) |
| *hst4*Δ | 5.31 (4.29-6.34) | 0.36 (0.29-0.43) | 15 (11-19) |
| *hst3*Δ *hst4*Δ | 11.3 (8.66-14.8) | 4.53 (3.79-5.42) | 2.5 (1.8-3.5) |
| *hst1*Δ | 1.34 (1.13-1.55) | 0.047 (0.033-0.063) | 29 (20-41) |
| *sir2*Δ | 2.12 (1.93-2.31) | 0.050 (0.040-0.059) | 43 (35-53) |
| *rpd3*Δ | 0.25 (0.22-0.29) | 0.026 (0.020-0.033) | 9.7 (7.4-13) |
| *gcn5*Δ | 0.98 (0.86-1.10) | 0.068 (0.054-0.082) | 15 (11-19) |
| *sir2*Δ *cac1*Δ | 4.44 (3.77-5.11) | 0.030 (0.015-0.046) | 151 (87-264) |
| *sir2*Δ *rtt106*Δ | 5.40 (4.36-6.43) | 0.12 (0.089-0.16) | 44 (31-62) |
| *sir2*Δ *cac1*Δ *rtt106*Δ | 6.32 (5.26-7.37) | 0.18 (0.13-0.24) | 35 (25-50) |
| *msh2*Δ *cac1*Δ | 3.18 (2.58-3.78) | 5.61 (5.01-6.20) | 0.57 (0.45-0.70) |
| *msh2*Δ *rtt106*Δ | 4.07 (3.49-4.65) | 3.35 (2.98-3.67) | 1.2 (1.0-1.4) |
| *msh2*Δ *sir2*Δ | 4.87 (4.23-5.50) | 3.58 (3.05-4.11) | 1.4 (1.1-1.7) |
| wild-type-2μ empty vector<sup>#</sup> | 0.74 (0.63-0.85) | 0.034 (0.022-0.048) | 22 (15-33) |
| wild-type-2μ MSH2-MSH6<sup>#</sup> | 5.69 (4.82-6.54) | 0.061 (0.043-0.080) | 94 (67-132) |
| *sir2*Δ-2μ empty vector | 4.35 (3.58-5.13) | 0.067 (0.045-0.089) | 66 (45-96) |
| *sir2*Δ-2μ MSH2-MSH6 | 4.55 (3.52-5.59) | 0.040 (0.027-0.053) | 116 (77-173) |
| *rtt106*Δ, 2μ empty vector | 2.01 (1.59-2.43) | 0.044 (0.028-0.061) | 46 (30-71) |
| *rtt106*Δ, 2μ MSH2-MSH6 | 2.49 (2.01-2.97) | 0.15 (0.10-0.20) | 17 (12-25) |

| | Cβ2/Cβ2-4L rate ($\times 10^{-6}$) | Cβ2/Cβ2 rate/<br>Cβ2/Cβ2-4L rate |
| --- | --- | --- |
| wild-type | 0.057 (0.042-0.072) | 9.2 (7.0-12.1) |
| *cac1*Δ | 0.070 (0.048-0.093) | 17 (12-25) |
| *asf1*Δ | 0.61 (0.48-0.74) | 6.5 (4.9-8.6) |
| *cac1*Δ *asf1*Δ | 0.25 (0.17-0.32) | 2.4 (1.4-4.3) |
| *sir2*Δ | 0.15 (0.12-0.18) | 14 (11-17) |

The mutations comprising the *pol30-8 allele* (R61A, D63A) were introduced into pEAA578 (*POL30∷KANMX*) by Q5 site-directed mutagenesis (New England Biolabs) to create the single step integrating vector pEAA633 (*pol30-8∷KANMX*). Plasmids pRS426 (*2μ, URA3*; Christianson *et al.* 1992) and pEAM272 (*MSH2, MSH6, 2μ, URA3;* Chakraborty, Dinh and Alani 2018) were used in the MSH overexpression experiments presented in Table 2.

### Inverted repeat recombination assay

Strains used to measure homologous and divergent recombination are listed in Table 1. Strains lacking plasmids were initially struck onto synthetic complete plates and those containing *2μ* plasmids were struck onto minimal dropout media plates (Rose, Winston, and Hieter 1990). A total of 10-117 single colonies per strain were then inoculated into 5 ml of synthetic complete or minimal dropout medium containing 4% galactose and 2% glycerol and grown to saturation for ∼2 days at 30°C. Appropriate dilutions of cells were plated onto minimal media (2% galactose, 2% glycerol) plates lacking histidine and the amino acid required to maintain the *2μ* plasmid (selective) and onto minimal media (2% glucose) plates lacking the amino acid required to maintain the *2μ* plasmid (permissive). Plates were incubated for 4 days at 30°C (5 days for experiments involving *rpd3Δ*) and then scored for frequency of His^+^ colonies. Rates of homologous and divergent recombination were calculated as described below.

### Analysis of mutation rates presented in Table 2

Asteris and Sarkar (1996) showed that Bayesian estimators of mutation rates from fluctuation experiments are more accurate than even the maximum likelihood estimator. We used an extension of their approach to calculate the posterior distribution of the mutation rate per cell division, *μ = m/N*_*t*_, where m is the estimated number of mutations and *N*_*t*_ is the total number of cells in the culture. This permitted credible intervals of the mutation rate ratios to be accurately calculated. Posterior distributions were calculated using the MSS (<u>M</u>a-<u>S</u>andri-<u>S</u>arkar) likelihood function (Ma, Sandri and Sarkar, 1992) for the Lea-Coulson model (Lea and Coulson 1949) with a non-informative (constant) prior over log *μ*. The results were insensitive to the choice of prior: Changing to a constant prior over *μ* or to a 1/*μ*^2^ prior changed the estimated mutation rate ratios by <10% of the distance to the credible interval boundaries (RMS deviation = 4%).

Depending on the mutation rates estimated in pilot experiments, fractions *f*, ranging between 0.005 to 0.4 of the cultures, were plated on non-permissive medium and counted. The total number of cells in the culture, *N*_*t*_, was measured in parallel in each case by plating a small aliquot on permissive medium. The likelihood function was calculated independently for each culture using the number of His^+^ colonies, *N*_*t*_, and *f*. The statistical error introduced by the partial plating with fraction *f* was included using the method of Zheng (2008); the statistical error in determining each *N*_*t*_ was included assuming Poisson sampling.

The variation of mutation rates between transformants of the same genotype, except for *hst3Δ hst4Δ*, were modest (Table S1). Therefore, the data for these genotypes were pooled and the mutation rates (Table 2 and Figure 3) were computed from the overall posterior distributions *p*(log *μ*_H_) and *p*(log *μ*_D_) using a quadratic loss function. The 95% credible intervals were computed using the highest posterior density. Figure 2 displays the geometric means of the homologous recombination (H) rate/divergent recombination (D) rate ratios computed using the marginal posterior distribution of *p*(log *μ*_H_ – log *μ*_D_), which was computed by integration over the joint distribution *p*(log *μ*_H_) × *p*(log *μ*_D_) (Gelman *et al.* 2014). The corresponding 95% credible intervals for log (*μ*_H_ /*μ*_*D*_) were computed from the highest posterior density over *p*(log *μ*_H_ – log *μ*_D_). The posterior distributions were very close to normal; the Hellinger distances between them and the best-fit normal distributions were 0.04; therefore, integrals were approximated using this assumption.

**Figure 1.**
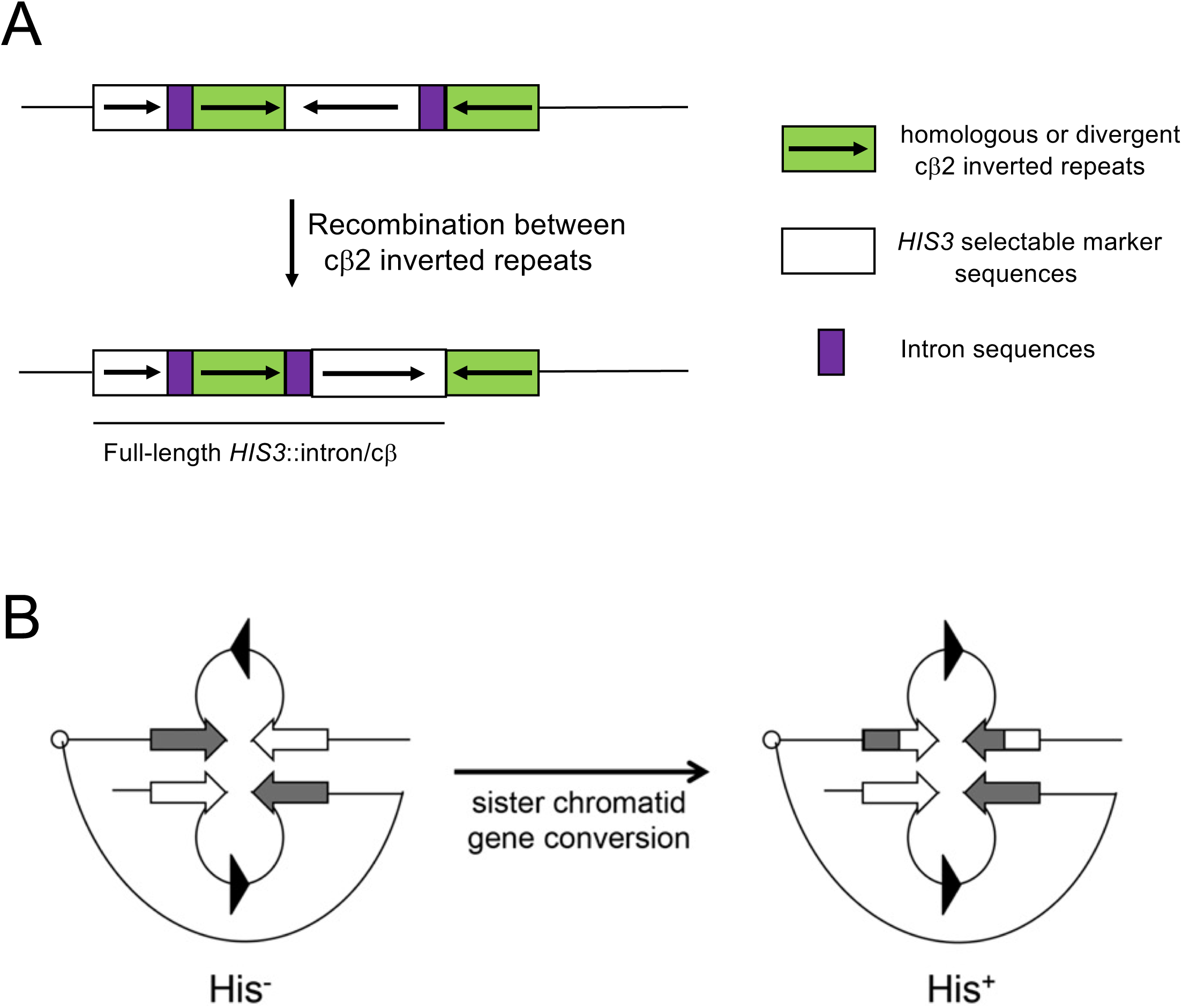
Schematic of an intron-based recombination assay involving inverted repeat sequences that form a functional *HIS3* reporter following HR (adapted from Nicholson *et al*. 2000). A. Repeat sequences depicted by the green boxes are either identical in sequence (homologous substrate) or differ (divergent) by four single nucleotide polymorphisms (cβ2/cβ2-ns) or four 4-nt insertions (cβ2/cβ2-4L). The cβ2/cβ2-ns and cβ2/cβ2-4L substrates are predicted to form base-base and 4-nt loop mismatches in heteroduplex DNA, respectively. In this assay, spontaneous His^+^ colonies result from recombination between the 350 bp repeat sequences that reorient *HIS3* (white boxes) and intron (purple boxes) sequences to yield a full-length *HIS3*∷intron gene. B. A model of how His^+^ recombinants arise in this system (Chen and Jinks-Robertson 1998). His^+^ recombinants are thought to result from gene conversion events between inverted repeat sequences present on sister chromatids (in this example, white and black arrows represent inverted repeat sequences) that reorient *HIS3* and intron sequences (triangles illustrate orientation of the *HIS3* and intron sequences). Homologous and divergent recombination rates for base–base and 4-nt loop substrates were calculated as described in the Materials and Methods, and the ratio of homologous to divergent recombination rates is presented as a measure of heteroduplex rejection efficiency.

**Figure 2.**
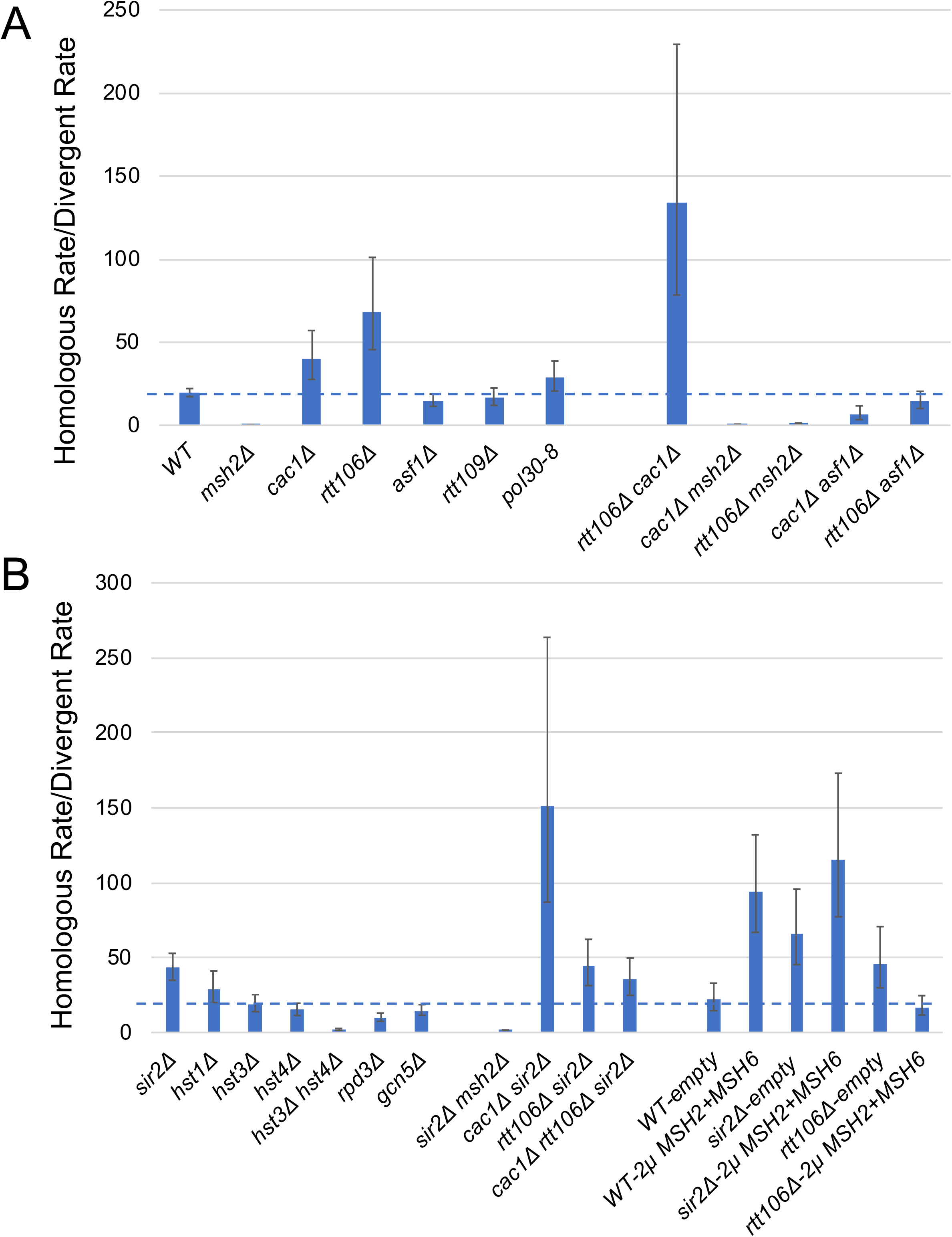
Recombination rates as measured in the inverted repeat reporter assay. Homologous and divergent recombination rates (calculated as described in the Materials and Methods, with 95% credible intervals, see Table 2) are shown in a bargraph for base-base mismatches for wild-type and the indicated mutant strains. A. Analysis of the effect of mutations in chromatin remodelers on the homologous rate/divergent rate ratio. B. Analysis of the effect of mutations in histone acetylases and deacetylases on the homologous rate/divergent rate ratio.

**Figure 3.**
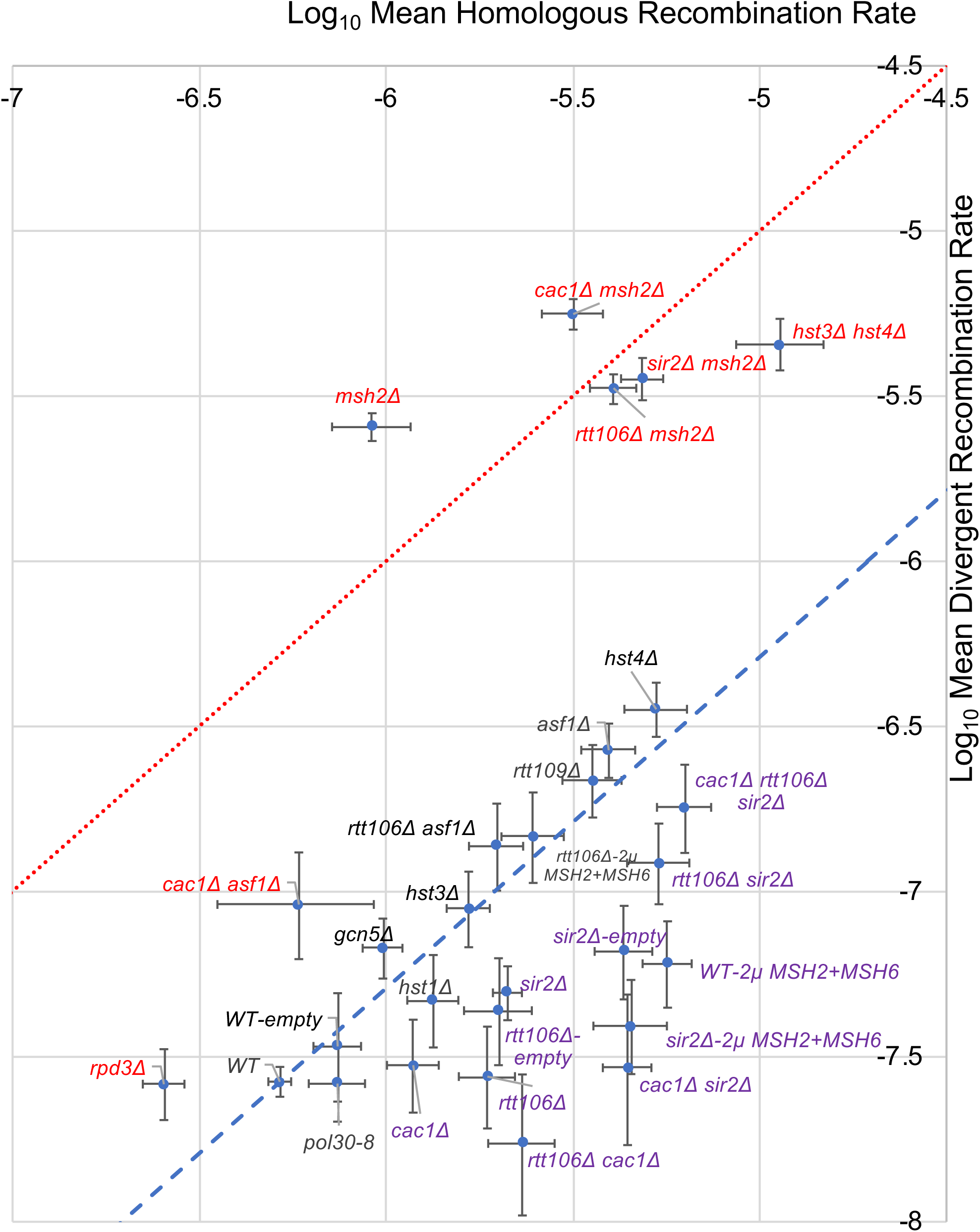
Roles for histone chaperones, acetylases and deacetylases in regulating heteroduplex rejection efficiency. A. Homologous and divergent recombination rates (calculated as described in the Materials and Methods) are shown for base-base mismatches for wild-type and the mutant strains presented in this study. The data in Table 2 were plotted to represent the least squared mean of the log_10_ transformed recombination rates (Materials and Methods). Error bars represent the 95% credible interval. The red dashed line represents a hypothetical homologous/divergent recombination ratio of 1, as seen in mutants defective in heteroduplex rejection. The blue dashed line represents a homologous/divergent recombination rate of 20, as seen in wild-type. Mutants that deviate from wild-type are shown with red fonts indicating a reduced heteroduplex rejection ratio and purple fonts indicating an increased ratio.

The *hst3Δ hst4Δ* data displayed anomalies that required special treatment. About 17% of the histidine dropout plates in which independent cultures of *hst3Δ hst4Δ* mutants were plated (both homologous and divergent strains) had zero colonies, or in one case, one colony, that were extremely inconsistent (p< 10^−6^) with the Lea-Coulson model (Lea and Coulson 1949) that provides the basis for the analysis. (The median number of colonies in the other cultures were > 100.) They were omitted since these outliers (8 homologous, 7 divergent independent cultures) would distort the analysis and yield extreme recombination rate outliers. This strain displays extremely high levels of genetic instability (Kadyrova *et al*. 2013), and thus the lack of His^+^ colonies on these plates likely resulted from this phenotype. Even when these values were included, the ratio of homologous recombination rate/divergent recombination rate increased by less than two-fold, presumably because the events that caused these unusual values were similar in the two strains.

Because of the above issue and possibly related two-fold variation in mutation rates of the different homologous transformants, we computed the mutation rates for each *hst3Δ hst4Δ* transformant separately. The weighted (inverse-variance) means and standard deviations of the transformant log *μ* values were computed. The ratio and 95% credible intervals were computed using the normal approximation to the posterior distribution.

### Repetition of experiments

The inverted repeat recombination assays were repeated on 2-4 days, with roughly an equal number of repetitions per day, and 2-4 independent transformants were analyzed for each genotype.

### Data availability

Strains and plasmids are available upon request. The authors affirm that all data necessary for confirming the conclusions of the article are present within the article, figures, and tables. Table S1, Variation in recombination rate between transformants, can be found in the GSA Figshare portal.

## Results

### Rationale for the experiments performed in this study

CAF-1 was shown to stabilize D-loops in *S. pombe* that occur as the result of template switching at replication forks (Pietrobon *et al*. 2014). This observation encouraged us to test if the histone chaperones CAF-1 and Rtt106 play roles in stabilizing recombination intermediates during HR, and in turn suppressing rejection. If these factors play such a role, heteroduplex rejection would occur more frequently in *cac1Δ* (deletion of the large subunit of CAF-1) and *rtt106Δ* mutants; however, heteroduplex rejection functions in these mutant backgrounds should remain dependent on mismatch recognition functions if CAF-1 and Rtt106 affect the stabilization of recombination intermediates, but not the mechanisms of anti-recombination mediated by heteroduplex rejection factors through mismatch recognition.

We also tested roles for other histone chaperones and chromatin modifying enzymes associated with DNA replication. These include Asf1, which binds to newly synthesized H3/H4 dimers that are then acetylated at H3K56 by Rtt109 (Tyler *et al.* 1999; Driscoll, Hudson and Jackson 2007; Han *et al*. 2007), and Rtt109, which mediates histone modification that promotes binding of H3 to the histone chaperones CAF-1 and Rtt106, and subsequently, the binding of CAF-1 to PCNA, facilitating histone deposition near the replication fork (Li *et al*. 2008). We were encouraged to study the above factors because much is known about the interactions and crosstalk between the replication apparatus and nucleosome assembly machinery. For example, CAF-1 interacts with the replication processivity clamp PCNA and Asf1 interacts with RFC, which loads PCNA, and the MCM helicase (Groth *et al*. 2007; Franco *et al.* 2005). In addition to their role in nucleosome assembly, Asf1 and Rtt109 indirectly promote nucleosome disassembly through H3K56 acetylation (Adkins, Howar and Tyler 2004; Adkins and Tyler 2004; Schwabish and Struhl 2006; Korber *et al.* 2006). The multifunctional roles of Asf1 and Rtt1109 (promoting nucleosome disassembly as well as binding of H3 to CAF-1 and Rtt106) thus make it difficult to predict what roles these factors might play in modulating heteroduplex rejection.

Finally, we were encouraged to test roles for the silencing factor Sir2 and related deacetylases (additional rationale provided below) based on work studies showing that Sir2 is recruited by CAF-1 and Rtt106 to sites of chromatin formation (Huang *et al*. 2007). In the model outlined above, histone chaperones would act in steps that involve the stabilization of recombination intermediates; Sir2 would act in conjunction with CAF-1 and Rtt106 to stabilize recombination intermediates. Thus, Sir2 would be predicted to provide additional steps to stabilize recombination intermediates, and *sir2Δ* mutants would be predicted to increase the frequency of heteroduplex rejection, perhaps to levels similar to that seen in *cac1Δ* and *rtt106Δ* mutants.

We used an inverted repeat recombination assay in baker’s yeast to study roles for chromatin modifiers in regulating the repair/rejection decision (Nicholson *et al*. 2000; Figure 1). Specifically, this assay measures spontaneous recombination events between homologous or divergent repeat sequences that reorient *HIS3* and intron sequences to yield a functional *HIS3* gene. Such events, thought to be initiated by DNA lesions that occur during or shortly after the replication of the recombination substrates, are consistent with repair through sister chromatid gene conversion (Chen and Jinks-Robertson 1998). For each genotype, we calculate the ratio of recombination rates (+/- 95% credible intervals) in a strain containing identical inverted repeat DNA substrates to those seen in a strain containing divergent inverted repeats (Materials and Methods). In the divergent strains heteroduplex recombination intermediates would form four single nucleotide mismatches or four 4-nucleotide loop mismatches in a 350 bp substrate (Figure 1). In wild-type this ratio is ∼20 for single nucleotide mismatches; in strains containing deletions in *MSH2* or *SGS1* the ratio ∼1, indicating that homologous and divergent recombination rates are similar, whereas deletions in *PMS1* and *MLH1* confer more modest effects (Table 2; Figure 2; Nicholson *et al.* 2000; Myung *et al*. 2001; Spell and Jinks-Robertson 2004). Similar genetic dependencies were seen in other assays that measure recombination between divergent DNA substrates (Selva *et al*. 1995; Sugawara *et al*. 2004). These observations have led to heteroduplex rejection models in which MSH proteins recognize mismatches in heteroduplex DNA and recruit the Sgs1-Top3-Rmi1 complex to unwind the recombination intermediate (Myung *et al*. 2001; Sugawara *et al*. 2004; Spell and Jinks-Robertson 2004).

### The absence of histone chaperones CAF-1 and Rtt106, but not Asf1 and Rtt109, improves anti-recombination

As shown in Table 2 and Figure 2, the *cac1Δ* mutation (deletion of the large subunit of CAF-1) increased the rejection ratio by two-fold, compared to wild-type, in recombination assays involving base-base or 4-nt loop mismatches. The *rtt106Δ* mutation conferred a three-to-four-fold increase in this ratio as measured in the base-base mismatch recombination assay. *msh2Δ cac1Δ* and *msh2Δ rtt106Δ* mutants showed rejection ratios similar to *msh2Δ* strains, indicating that the increases in the rejection ratio in *cac1Δ* and *rtt106Δ* strains were dependent on the rejection machinery. Interestingly, the *cac1Δ rtt106Δ* double mutant showed a rejection ratio significantly higher (seven-fold increase compared to wild-type) than the single mutants, suggesting a redundant function for these two factors analogous to redundant roles for these proteins in DNA replication (Li *et al*. 2008). Together, these observations suggested that deposition of histones by CAF-1 and Rtt106 during double-strand break repair stabilized recombination intermediates, making them less accessible to heteroduplex rejection factors.

In contrast, deleting Asf1 or Rtt109, factors thought to act upstream of CAF-1 and Rtt106, did not confer an effect on heteroduplex rejection (Table 2). These factors are required to acetylate Histone H3 at K56, and the resulting H3K56ac-H4 dimers are then transferred to histone chaperones such as CAF-1 and Rtt106 (Schneider *et al*. 2006; Han *et al*. 2007; Driscoll, Hudson and Jackson 2007; Tsubota *et al*. 2007). Interestingly, the *asf1Δ* mutation suppressed the increase in rejection seen in *cac1Δ* or *rtt106Δ* mutants, and for *cac1Δ asf1Δ* mutants the rejection ratio was lower than wild-type. Additionally, we performed recombination assays in 4-nt loop mismatch strains where heteroduplex rejection depends primarily on the Msh2-Msh3-complex (Nicholson *et al*. 2000). We found that the *asf1Δ* mutation had minimal if any effect on rejection, and that it suppressed the increased rejection seen in *cac1Δ* strains (to lower than wild-type), in agreement with observations from base-base mismatch strains (Table 2; Figure 2). This observation also indicates that the effects on heteroduplex rejection seen in chromatin modifier mutants in the base pair mismatch strains (where heteroduplex rejection is primarily dependent on Msh2-Msh6) were not specific to a particular MSH complex.

Previous studies showed that mutants defective in the CAF-1 complex are sensitive to DNA damaging agents, and have suggested that during DNA repair CAF-1 is recruited by the replication processivity clamp PCNA to facilitate DNA synthesis repair steps (Moggs *et al.* 2000; Linger and Tyler 2005). The Stillman group identified in baker’s yeast an allele of *POL30* (gene encoding PCNA; *pol30-8* (R61A, D63A)) that displayed reduced binding to CAF-1 and compromised recruitment of CAF-1 to replicating DNA (Shibahara and Stillman 1999; Zhang, Shibahara, and Stillman 2000), and Linger and Tyler (2005) showed that *pol30-8* mutant yeasts were similarly sensitive to DNA damaging agents as *cac2Δ* (subunit of the heterotrimeric CAF-1 complex) and *cac2Δ pol30-8* mutants. As shown in Table 2 and Figure 2, the *pol30-8* allele, which conferred sensitivity to MMS, increased heteroduplex rejection by slightly (1.4-fold), although the 95% credible interval of the ratio is large (0.8 to 2.7), indicating a significant overlap between the rejection ratios for wild-type and *pol30-8*.

### Deletion of the Sir2 silencing factor increases anti-recombination

The silencing factors Sir2 and Sir3 are recruited by the histone chaperones CAF-1 and Rtt106 to sites of heterochromatin formation. In the absence of CAF-1 and Rtt106, Sir proteins are mis-localized (Huang *et al*. 2007). These observations encouraged us to test if the histone deacetylases Hst1, Hst3, Hst4 or Sir2 are involved in the regulation of rejection efficiency. We were further encouraged to test these factors because Tamburini and Tyler (2005) showed in *S. cerevisiae* that the histone acetylases Gcn5 and Esa1 (essential for viability) and the histone deacetylases Rpd3, Sir2 and Hst1 were recruited to an HO endonuclease induced lesion during HR. We were also interested in Hst3 and Hst4, which act to remove H3K56ac marks from newly generated chromatin in G2/M, because studies from Munoz-Galvan *et al*. (2013) suggested that acetylation and deacetylation of H3K56 were important for selecting the sister chromatid as a template for repair of DSBs that occur during DNA replication. As shown in Table 2 and Figure 2, the *sir2Δ* mutation increased the efficiency of heteroduplex rejection by 2.2 (1.7 to 2.8, 95% credible intervals)- and 1.5-fold (1.1 to 2.1, 95% credible intervals) in the nucleotide substitution and 4-nt loop strains, respectively. The *sir2Δ msh2Δ* mutant displayed a homoduplex/divergent recombination ratio similar to *msh2*Δ, indicating that the increased rejection ratios seen in *sir2*Δ strains required the heteroduplex rejection machinery. Compared to wild-type, deletion of *HST3* or *HST4* conferred no significant effect on the rejection ratio. However, the *hst3Δ hst4Δ* double mutant showed an 8.0-fold *reduction* in anti-recombination of base-base mismatches compared to wild-type strains, but the strain also showed variation in recombination rates between transformants (unrelated to whether the strain contains the homologous or divergent recombination substrate; see Materials and Methods) that is likely due to the high genomic instability (Kadyrova *et al.* 2013) seen in this mutant. Thus, there is a need for caution in interpreting the *hst3Δ hst4Δ* results.

The increased heteroduplex rejection phenotype observed in *sir2Δ* strains and the opposite phenotype seen in *hst3Δ hst4Δ* strains encouraged us to test if deletion mutations in the histone deacetylase Hst1, the acetylase Gcn5, and Rpd3, a modifier of Sir2 function, affected rejection. As shown in Table 2 and Figure 2, *hst1Δ* and *gcn5Δ* mutations did not confer an effect. We then tested a strain deleted for Rpd3, which encodes a subunit of both the Rdp3S and Rpd3L histone deacetylase complexes that act in chromatin remodeling. Several studies reported that Rpd3 and Sir2 have antagonistic effects on silent chromatin propagation and replication timing (Zhou *et al*. 2009; Thurtle-Schmidt, Dodson, and Rine 2016; Yoshida *et al*. 2014; Ehrentraut *et al*. 2010). The *rpd3Δ* mutation decreased rejection two-fold in the nucleotide substitution strain. Together these observations provide evidence that chromatin modifiers can suppress and enhance heteroduplex rejection (see Discussion).

### *SIR2, CAC1,* and *RTT106* may act in similar steps

To determine if CAF-1 and Rtt106 function in similar steps as Sir2 to prevent heteroduplex rejection, we examine the phenotype of double mutants. *rtt106Δ sir2Δ* mutant strains exhibited rejection ratios similar to those of *rtt106Δ* or *sir2Δ* single mutants where as *cac1Δ sir2Δ* showed a higher ratio compared to *cac1Δ* or *sir2Δ* (Table 2 and Figure 2). Curiously, the rejection ratio was lower or similar for *sir2Δ cac1Δ rtt106Δ* mutants compared to any of the single mutants, indicating a more complex genetic interaction (see below).

Previously we showed that co-overexpression of the Msh2 and Msh6 MMR proteins in wild-type strains increased the frequency of heteroduplex rejection in the inverted repeat assay by 3.5-fold, and hypothesized that this was due to the increased availability of functional MSH complexes acting in heteroduplex rejection (Chakraborty *et al*. 2016; 2018). Because both the *sir2Δ* mutation and Msh2 and Msh6 co-overexpression increased rejection, we asked if the increased rejection seen in *sir2Δ* strains would rise to an even higher level in the presence of pEAM272 (*2*μ*, MSH2, MSH6* plasmid). Such an experiment tests if the increases in rejection seen in the two conditions reflect common or distinct regulatory steps. In wild-type strains bearing pEAM272 the Msh2-Msh6 complex is overexpressed by ∼8-fold (Chakraborty, Dinh and Alani 2018). Wild-type strains containing pEAM272 display a 4.3-fold increase, compared to those containing the empty vector pRS426, in the rejection ratio, consistent with an increased concentration of Msh2-Msh6 resulting in improved rejection by increasing the likelihood of mismatch recognition in heteroduplex DNA (Table 2). *sir2Δ* strains containing pEAM272 display a higher rejection ratio (1.8-fold (1.1 to 3.1, 95% credible intervals) higher than *sir2Δ* with empty vector, but with some overlap in 95% credible intervals), suggesting that the *SIR2* effect on anti-recombination may not be similar to that seen in the wild-type strain containing pEAM272. One interpretation of this observation is that the Sir2 effect on rejection occurs in steps that compete with mismatch recognition. Curiously, *rtt106Δ* strains containing pEAM272 showed a rejection ratio that was similar to the wild-type lacking pEAM272 (Table 2), indicating a more complex phenotype reminiscent of the decreased rejection ratio seen in *sir2Δ cac1Δ rtt106Δ* triple mutants.

### Altered recombination rates in chromatin remodeling mutants do not correlate with their heteroduplex rejection phenotypes

The inverted repeat assay detects spontaneous recombination events. *msh2Δ* mutants, which are defective in heteroduplex rejection, display elevated levels of homologous recombination (Table 2; Datta *et al.* 1997; Nicholson *et al*. 2000). Previously the Jinks-Robertson group hypothesized that the increased HR seen in *msh2Δ* reflects the fact that the length of perfect homology required to avoid heteroduplex rejection (610 bp) is larger than the length of the 350 bp repeats present in the inverted repeat substrate (Datta *et al.* 1997). However, for other mutants, increased HR could be due to increases in the formation of DNA lesions that increase the initiation of such events. As shown in Figure 3 and Table 2, the rates of recombination between homologous sequences in the mutants analyzed in this study vary, but do not appear to correlate to changes in their repair/rejection ratios. For example, *asf1Δ* and *rtt109Δ* mutants show recombination levels higher than *msh2Δ*, but a homologous/divergent recombination ratio similar to wild-type. In contrast, *cac1Δ* and *rtt106Δ*, which show homologous recombination levels between wild-type and *msh2Δ*, and *sir2Δ*, which shows recombination levels similar to *msh2Δ*, displayed homologous/divergent recombination ratios higher than wild-type. A similar lack of correlation was seen for the double mutant combinations presented. This information suggests that overall levels of homologous recombination do not impact the rejection ratio.

## Discussion

This study focused on understanding roles for chromatin structure and modifications in regulating the heteroduplex rejection/DNA repair decision. Improving repair at the cost of reduced fidelity can lead to gene conversion, chromosomal rearrangement and loss of heterozygosity, whereas high fidelity can compromise repair efficiency (reviewed in Chakraborty and Alani 2016). We found that the histone chaperones CAF-1 and Rtt106 and the Sir2 deacetylase act to suppress heteroduplex rejection. In contrast, a large set of factors involved in nucleosome assembly during DNA replication or in modifying histones (Rtt109, Hst1, Gcn5) do not affect rejection efficiency. These results are consistent with repair pathways in which the presence of nucleosomes at DNA lesions acts to stabilize recombination intermediates and inhibit anti-recombination (Figure 4).

**Figure 4.**
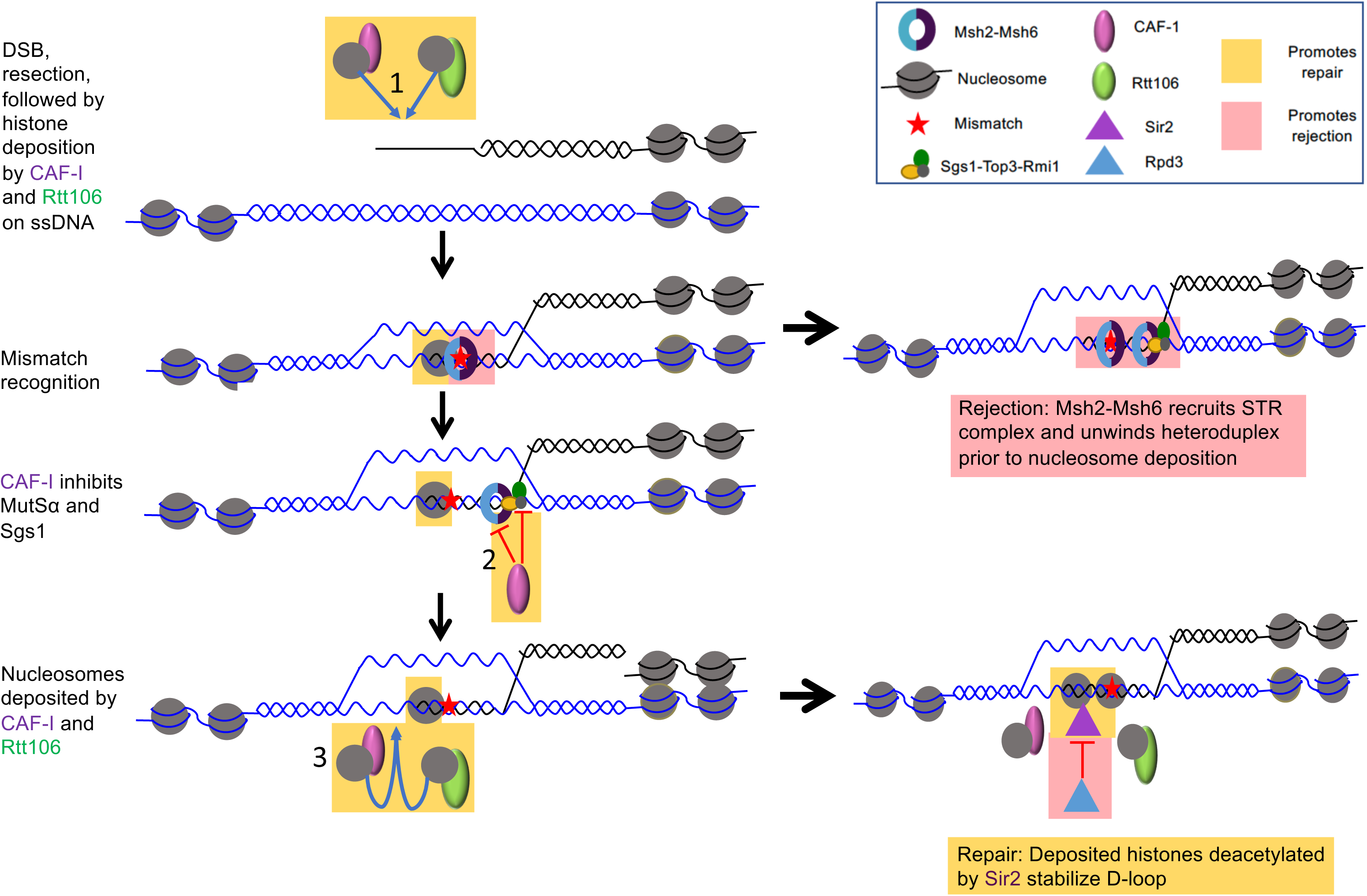
A model outlining possible steps during homologous recombination where chromatin modifiers regulate the repair/rejection decision. We hypothesize that factors promoting nucleosome assembly/maintenance such as CAF-1 and Rtt106 deposit nucleosomes at DNA lesions independent of DNA synthesis on ssDNA prior to strand invasion (1). Such deposition/maintenance, coupled with the formation of Sir2-dependent chromatin marks that promote a closed chromatin state, stabilizes D-loops to suppress rejection. Additionally, we hypothesize that CAF-1 physically interacts with Msh6 and possibly Sgs1 to inhibit the rejection factors from unwinding heteroduplex substrates (2). Finally, DNA synthesis coupled nucleosome deposition by CAF-1 and Rtt106 combined with Sir2 dependent modifications of these nucleosomes (negatively regulated by Rpd3) are likely to stabilize heteroduplex D-loop intermediates, making it more difficult for rejection factors to unwind the intermediates (3). In this model MSH proteins are recruited to a DSB either directly or through specific histone marks, and nucleosome maintenance limits the time frame in which anti-recombination can be performed and promotes repair of the broken chromosome and ultimately mismatch repair steps (see text for additional details). Factors/steps that promote rejection are highlighted in pink boxes and those that promote repair are highlighted in yellow boxes.

The model presented in Figure 4 outlines a recombination event involving divergent sequences initiated by a DSB. DNA mismatches in heteroduplex DNA that form during strand invasion are recognized by MMR proteins which in turn recruit the Sgs1-Top3-Rmi1 helicase-topoisomerase complex to unwind and reject the recombination intermediate. In this model, heteroduplex rejection is repressed by the presence of nucleosomes in a process regulated by CAF-1 and Rtt106, representing a regulatory step in the repair/rejection decision. The maintenance of nucleosomes by the histone chaperones CAF-1 and Rtt106 acts to stabilize heteroduplex DNA. This is followed by Sir2 deacetylating newly deposited histones, creating a “closed” chromatin structure that prevents rejection but promotes the completion of repair through DNA synthesis steps (see the access-repair-restore model of Tamburini and Tyler 2005). In other words, a closed chromatin structure, particularly in later stages in repair, would promote completion of homologous recombination, and prevent access to MMR factors that can act in heteroduplex rejection. Thus, the window for heteroduplex rejection would occur prior to the formation of closed chromatin.

The model presented in Figure 4 is supported by the following observations:

1. Deleting *CAC1* or *RTT106* resulted in increased heteroduplex rejection. Furthermore, *cac1Δ rtt106Δ* strains showed an even higher increase in heteroduplex rejection, consistent with CAF-1 and Rtt106 acting redundantly in nucleosome assembly (Table 2; Li *et al*. 2008).
2. A number of studies have shown that MMR and nucleosome assembly during replication are mutually inhibitory processes and that human MutSα interacts physically with CAF-1 (Li *et al*. 2009; Kadyrova, Blanko and Kadyrov 2011; Schopf *et al.* 2012; Blanko, Kadyrova and Kadyrov 2016). Thus, in a manner analogous to their interaction during post-replicative MMR, nucleosome deposition could suppress MMR mediated heteroduplex rejection, forcing the latter process to occur in the time window after strand invasion and before nucleosome deposition.
3. Various studies have shown in fission yeast and humans that RecQ helicases (Rqh1 in *S. pombe* and BLM and WRN in humans) physically interact with the large subunit of CAF-1 (Pietrobon *et al*. 2014; Jiao *et al*. 2004; Jiao *et al*. 2007). Additionally, Pietrobon *et al*. (2014) showed that CAF-1 suppresses D-loop disassembly by Rqh1 during template switching. Based on these observations, it is possible that CAF-1 physically interacts with Sgs1 in budding yeast and counteracts its unwinding activity during heteroduplex rejection.
4. The removal of 3’ non-homologous tails during SSA acts as a temporal switch; rejection is favored before tail removal, prior to DNA synthesis steps, and repair, after (Chakraborty *et al*. 2016). Thus, analogous to 3’ non-homologous tail removal during SSA, nucleosome maintenance by CAF-1 and Rtt106, followed by nucleosome deacetylation by Sir2 during HR could provide another type of temporal commitment step that regulates the rejection vs repair decision. In support of this idea, Tamburini and Tyler (2005) showed that Sir2 localizes to sites of DSBs after histone acetylases such as Gcn5 and Esa1, suggesting that Sir2 is likely to localize to sites of HR at later stages of repair, likely after the deposition of nucleosomes, in order to modify histones by deacetylating them and further compacting the repair substrates making it harder for the rejection machinery to act.

Together, these observations support the idea that CAF-1 and Rtt106 function redundantly to deposit nucleosomes on recombination intermediates that stabilize the DNA heteroduplex and suppress rejection.

### *asf1Δ* mutation suppresses the hyper-rejection phenotype seen in *cac1Δ* and *rtt106Δ* strains

Curiously, the *asf1Δ* and *rtt109Δ* mutations did not alter heteroduplex rejection ratios, and *asf1Δ* suppressed the increased ratio seen in *cac1Δ* and *rtt106Δ* strains (both base-base and 4-nt loop mismatch substrates). This was surprising, given that Asf1 and Rtt109 act upstream of CAF-1 and Rtt106 in the nucleosome deposition pathway. Asf1 and Rtt109 have been implicated in nucleosome removal during DNA replication and transcriptional activation (Groth *et al*. 2007; Ransom, Dennehey, and Tyler 2010; Adkins, Howar and Tyler 2004; Adkins and Tyler 2004; Schwabish and Struhl 2006; Korber *et al.* 2006). Such a nucleosome removal activity could also act during DNA recombination, and in its absence, lead to chromatin acting to stabilize strand invasion intermediates that are refractory to heteroduplex rejection. In this model, a lack of, or delay in, nucleosome deposition in *cac1Δ* or *rtt106Δ* mutants resulting in increased rejection could be compensated for by the lack of nucleosome removal in *asf1Δ* mutants. Such a scenario is supported by recent *in vivo* and *in vitro* findings indicating that histones are present at ssDNA (Adkins *et al*. 2017; Huang *et al*. 2018). Adkins *et al*. (2017) also showed *in vitro* that histones remain bound to ssDNA as resection proceeds for longer distances.

We recognize that testing in the inverted repeat assay the effect of *H3K56Q* and *H3K56R* mutations, which mimic histone H3 acetylation and lack of acetylation at lysine 56, respectively (summarized in Kadyrova *et al*. 2013), could provide additional insights into the unexpected finding that *asf1Δ* suppresses the elevated rejection phenotype seen in *cac1Δ* and *rrt106Δ* mutants. However, H3K56 acetylation is required for both histone disassembly by Asf1, and for histone assembly by CAF-1 and Rtt106 (Tyler *et al*. 1999; Driscoll, Hudson and Jackson 2007; Han *et al.* 2007; Li *et al*. 2008; Williams, Truong and Tyler 2008). Thus, similar to what was seen in an *asf1* mutant, histone modifications would likely inhibit or promote (depending on the mutant) both histone disassembly and assembly and thus a test of histone mutant alleles in heteroduplex rejection assays may not provide additional insights. More detailed analyses are planned to understand the *asf1Δ* suppression phenotype.

### Does CAF-1 regulation of heteroduplex rejection depend on its interaction with PCNA?

CAF-1 has been implicated in DNA synthesis coupled nucleosome deposition via its interaction with PCNA (Shibahara and Stillman 1999; Zhang, Shibahara, and Stillman 2000; Krawitz, Kama, and Kaufman 2002). We found that strains bearing the *pol30-8* allele, which significantly weakens the PCNA-CAF-1 interaction, showed heteroduplex rejection ratios only slightly higher than wild-type (Table 2; Figure 2). One explanation for this phenotype is that CAF-1 regulation of heteroduplex rejection does not depend, or only partially depends, on its interaction with PCNA. In support of the former possibility, Hoek, Myers, and Stillman (2011) found that a mutation in human CAF-1 that causes a defect in PCNA interactions (N-terminal truncation of the p150 subunit of CAF-1) does not affect the recruitment of CAF-1 to sites of DNA damage or confer sensitivity to DNA damaging agents. They also showed direct interactions between CAF-1 and the KU complex and 14-3-3 proteins, both of which are involved in DNA damage responses. In addition, during baker’s yeast meiosis, CAF-1 is recruited independently of its interaction with PCNA to meiotic double-strand breaks at a step prior to strand invasion, though deletion of the large subunit of CAF-1 did not affect meiotic double-strand break repair or crossover formation (Brachet *et al*. 2015). Finally, Huang *et al*. (2018) have suggested that CAF-1 along with ASF1 are involved in chromatin assembly on ssDNA prior to Rad51 nucleoprotein filament formation, implying that it occurs in a PCNA independent manner, as PCNA might be expected to localize much later during DNA synthesis steps.

### MMR factors are likely to associate with sites of recombination early in the process

Our data suggest that the deposition of nucleosomes and their subsequent modification are likely to suppress rejection and limit the time window during which the rejection machinery can act to unwind recombination intermediates involving divergent substrates. Previously we showed that during SSA, removal of 3’ non-homologous tails serves a similar role with respect to providing a limited time frame for rejection to occur (Chakraborty *et al*. 2016). Several studies also support this idea. For example, Li *et al*. (2013) showed in mammalian cells that MSH2-MSH6 is recruited via a PWWP motif in MSH6 to H3K36me3 histones before or during early S-phase. In the context of rejection, this would ensure that mismatch recognition proteins are localized to chromatin before replication fork stalling related recombination events occur. Although yeast Msh6 and human MSH3 proteins do not have PWWP motifs, one can imagine that these proteins may be recruited by other histone marks to chromatin. Additionally, MSH proteins have been shown to localize rapidly to DSBs, even in the absence of a donor template, and act in the DNA damage response (Evans *et al*. 2000; Lyndaker, Goldfarb, and Alani 2008; Hong *et al*. 2008; Burdova *et al*. 2015). Interestingly, overexpression of Msh2 and Msh6 increased, albeit weakly, the rejection ratio further in *sir2Δ* strains, suggesting that mismatch recognition effects on rejection are distinct and are likely to precede changes in chromatin structure (Table 2). Taken together, these data suggest that mismatch recognition proteins localize to sites of recombination either before DSBs are formed or soon afterwards, and that the chromatin environment could alter this localization and thus impact heteroduplex rejection.

### Why do *rpd3Δ* strains display defects in heteroduplex rejection?

A large-scale screen for mutants defective in trinucleotide repeat instability identified a mutation in Sin3, which is a subunit of the histone deacetylases Rpd3L and Rpd3S (Debacker *et al*. 2012). In their analysis, *sin3* mutants displayed significant reductions (9 to 18-fold) in expansion rates for a trinucleotide repeat reporter. Trinucleotide repeat expansions are also suppressed by disruption of factors involved in heteroduplex rejection such as Msh2 and Msh3; such MSH factors are hypothesized to promote trinucleotide repeat expansions by stabilizing slipped strand intermediates or through altered MMR (McMurray 2010). Based on these observations, the defect in heteroduplex rejection observed in *rpd3Δ* strains could reflect the down-regulation of MSH factors that act in mismatch recognition. Alternatively, the *rpd3Δ* phenotype results from improved localization of Sir2 to an initiating DSB site, thus lowering rejection. The latter explanation fits with observations obtained from Zhou *et al*. (2009), who showed that in the absence of Rpd3, increased localization was seen for Sir2 at telomeres and homolgous mating type loci (HM), leading to an extension of silent chromatin in these areas.

### Can the heteroduplex rejection machinery be saturated?

As shown in Table 2 and Figure 2, the *hst3Δ hst4Δ* double mutant appears strongly compromised for anti-recombination and displays high rates of homologous recombination. Kadyrova *et al.* (2013) showed that *hst3Δ hst4Δ* mutants display very high mutation rates that result from base substitutions, 1-bp insertions/deletions, and spontaneous gross chromosomal rearrangements. The rate of mutations in *hst3Δ hst4Δ*, as measured in forward mutation and reversion assays, was similar to that seen in MMR defective strains, and *msh2Δ hst3Δ hst4Δ* triple mutants displayed greater than additive mutation rates compared to *msh2Δ* and *hst3Δ hst4Δ*. Based on these and other observations they proposed that Hst3 and Hst4 participate in genetic stability mechanisms that work with mismatch repair and replicative polymerase proofreading mechanisms to suppress spontaneous mutagenesis. In this framework, the decreased rejection seen in *hst3Δ hst4Δ* strains could result from high rates of mutagenesis saturating the MMR machinery and thus reducing the pool of MSH proteins available to participate in heteroduplex rejection. Another possibility is that the lack of Hst3 and Hst4 leads to higher levels of H3K56 acetylation, which in turn favors nucleosome deposition by CAF-1 and Rtt106 factors that selectively bind to H3K56 acetylated histones. Such a situation could stabilize the strand invasion intermediate and suppress anti-recombination. In contrast, the presence of Hst3 and Hst4 would promote rejection by deacetylating H3K56 and thus suppress nucleosome deposition by CAF-1 and Rtt106. In support of this idea, Celic *et al*. (2006) showed that H3K56 sites are hyperacetylated in yeast lacking Hst3 and Hst4. Additional studies will be required to distinguish between these models.

### Do chromatin modification factors have indirect effects on heteroduplex rejection?

Studies in yeast showed that the Fun30, RSC and INO80 chromatin remodelers promote resection of DNA DSB ends (Chen *et al*. 2012; Lademann *et al*. 2017; Daley *et al*. 2015), and studies in human cells showed that ASF1 protects such ends from excessive resection (Huang *et al*. 2018). Could changes in resection rates impact heteroduplex rejection? Resection of DNA DSB ends is a critical initiating step in homologous recombination because it generates 3’ single strand tails that participate in the formation of heteroduplex DNA. Thus, changes in resection rates could impact the stability of recombination intermediates by altering heteroduplex tract lengths. Consistent with this idea, a recent study in yeast showed that defects in resection as well as increased amounts of the single-strand binding protein RPA enhanced the efficiency of repair through an ectopic donor locus (Lee *et al*. 2016). We were unable to test effects on resection because His^+^ recombinants result from the repair of spontaneous DNA lesions. However, Datta *et al.* (1997) estimated in the system we used, the length of perfect homology required to initiate stable heteroduplex formation (20 bp), and avoid heteroduplex rejection (610 bp, which is larger than the 350 bp repeat). These estimates, the work of Lee *et al.* (2016), and our finding that homologous recombination is not decreased in the chromatin remodeling mutants analyzed, suggest to us that the *increased* rejection seen in *cac1Δ, rtt106Δ*, and *sir2Δ* mutants is not likely due to alterations in resection rates, though we cannot exclude the possibility that greater resection could lead to longer heteroduplex tract intermediates with more mismatches that are substrates for rejection. Also, we cannot exclude the possibility that the *decreased* rejection seen in *hst3Δ hst4Δ* and *rpd3Δ* mutants resulted from severely limited resection if only short heteroduplex tracts form that contain perfect homology tracts, without branch migration steps that would lead to the formation of mismatches.

We also recognize that the homologous/divergent recombination ratio cannot tell us if chromatin modifying mutations affect the utilization of specific recombination pathways that have different sensitivities to heteroduplex rejection. For example, Spell and Jinks-Robertson (2003; 2004) showed that Rad51-dependent recombination had more stringent requirements for homology between recombination substrates than Rad51-independent pathways. Using the same assay that we used here, they found that a null mutation in the Srs2 helicase, which acts to prevent Rad51-dependent recombination, conferred an increase in the homologous/divergent ratio, a phenotype similar to what we observed in *cac1Δ* and *rtt106Δ* mutants (Spell and Jinks-Robertson 2003; 2004; Krejci *et al*. 2003; Veaute *et al*. 2003). Based on these observations, Spell and Jinks-Robertson (2004) suggested that increased rejection occurred in *srs2Δ* because in such an environment Rad51-dependent recombination is favored, leading to more stringent homology requirements. Interestingly, they found that the increased rejection seen in *srs2Δ* mutants was suppressed by the *msh2Δ* mutation, suggesting that “much of the prevention of hyperrecombination between divergent recombination substrates is dependent on mismatch recognition, which also would be expected for *RAD51*-dependent recombination.” We observed a similar suppression in *cac1Δ msh2Δ* and *rtt106Δ msh2Δ* mutants (Table 2), suggesting that even if different recombination pathways are utilized in *rtt106Δ* and *cac1Δ* strains, the heteroduplex rejection machinery in these strains responds to mismatch recognition. The above concerns will need to be further explored in recombination systems where heteroduplex rejection can be monitored through DSB events induced at specific sites (e.g. HO, I-*Sce*I).

## Acknowledgments

We thank members of the Alani Lab, Jessica Tyler, Mike Fasullo, Michael Lichten and Scott Keeney for helpful comments and advice, and Sue Jinks-Robertson for providing us with the inverted repeat recombination strains SJR769, GCY615 and GCY559. We are grateful to Deniz Akdemir, Cornell Statistical Consulting Unit for his advice on the statistical analysis. U.C., B.M. and E.A. were supported by National Institutes of Health (NIH) grant GM53085. The funders had no role in study design, data collection and analysis, decision to publish, or preparation of the manuscript. The content is solely the responsibility of the authors and does not necessarily represent the official views of the National Institute of General Medical Sciences or the National Institutes of Health.

---

**Table S1.**
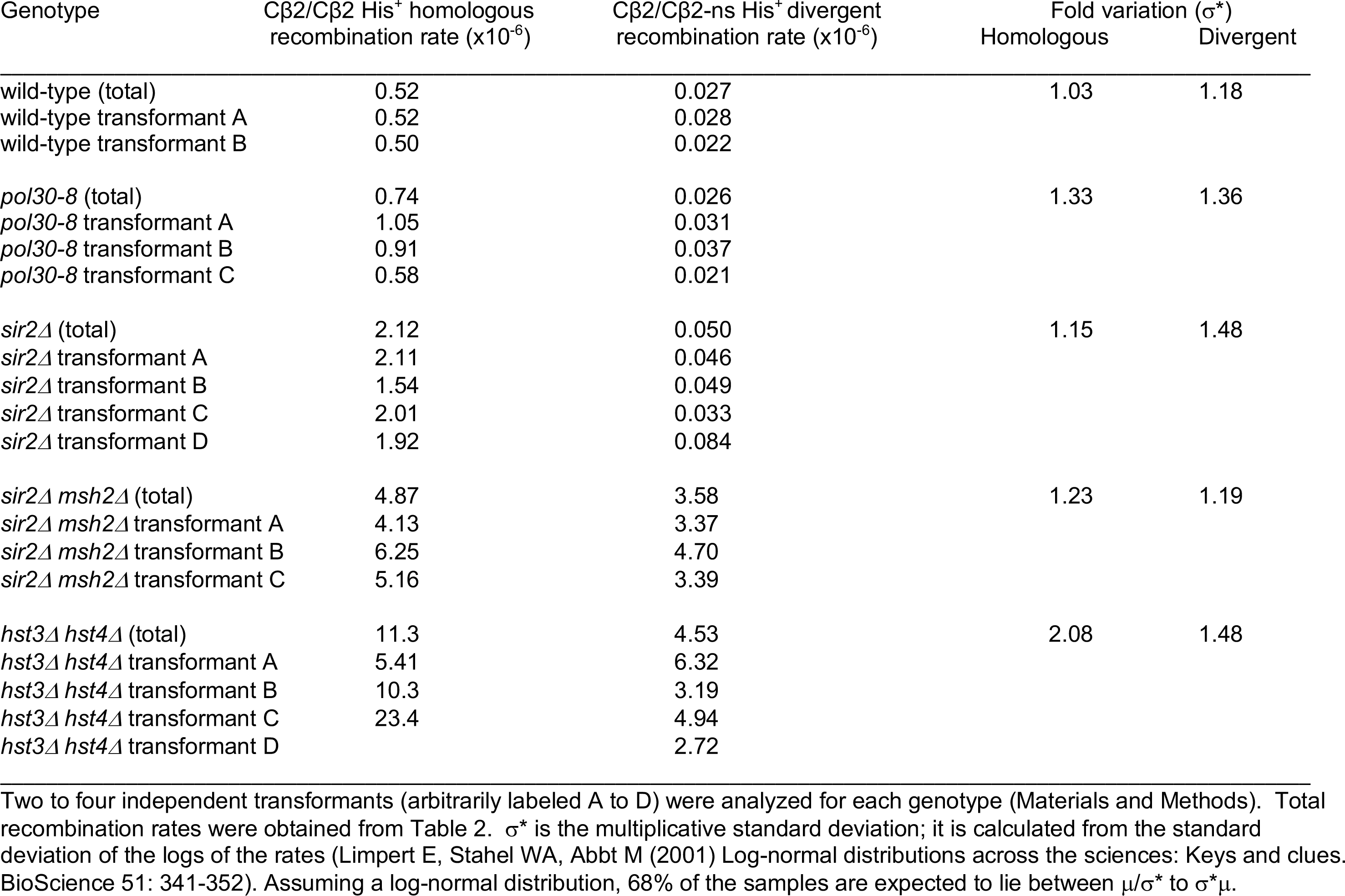
Variation in recombination rate between transformants.

